# Liquid foam therapy (LiFT) for homogenous distribution of exogenous pulmonary surfactant in ARDS

**DOI:** 10.1101/2020.11.30.403337

**Authors:** Rami Fishler, Yan Ostrovski, Avital Frenkel, Simon Dorfman, Vera Brod, Tali Haas, Dan Waisman, Josué Sznitman

**Author notes:** These authors contributed equally: Rami Fishler, Yan Ostrovski,Avital Frenkel.

## Abstract

Lung surfactant dysfunction has a critical role in the pathophysiology of acute respiratory distress syndrome (ARDS). Yet, efforts to treat ARDS patients with liquid instillations of exogenous surfactant have so far failed. One of the ongoing challenges in surfactant therapy is obtaining a homogeneous distribution of surfactant within the lungs despite an inherent tendency to non-uniform spreading, owing amongst others to the influence of gravity. Here, we show that liquid foam therapy (LiFT), where surfactant is foamed prior to intratracheal administration, may improve notably surfactant distribution while maintaining safety and efficacy. We first show quantitatively that a foamed surrogate surfactant solution distributes more uniformly in *ex vivo* pig lungs compared to endotracheal instillations of the liquid solution, while maintaining pulmonary airway pressures within a safe range. Next, we demonstrate that a foamed commercial surfactant preparation (Infasurf) is effective in an established *in vivo* rat lung lavage model of ARDS. Our results suggest that LiFT may be more effective than liquid instillations for treating ARDS and serve as a proof-of-principle towards large animal and clinical trials.

## introduction

Acute respiratory distress syndrome (ARDS) is a life-threatening inflammatory type of respiratory failure resulting for example from pneumonia, sepsis, drowning, and most recently the COVID-19 infection^1–3^. Among the central factors underlying the pathophysiology of ARDS lies critically the dysfunction of lung surfactant^4,5^. In turn, the administration of exogenous lung surfactant into the lungs of ARDS patients has been hypothesized as a promising therapeutic strategy with the potential to improve lung mechanics and oxygenation, while concurrently reducing ventilation-induced injury^4,6^. The use of intratracheal liquid instillations of lung surfactant is known to substantially reduce mortality in premature neonates suffering from respiratory distress syndrome (RDS) and has thus become an established rescue therapy in neonatal populations over the past decades^7,8^. In parallel, surfactant therapy using liquid instillations has shown promising results in medium-sized clinical trials of pediatric and adult ARDS patients^9,10^. Yet, larger clinical studies in ARDS patients have consistently failed to show clinical benefits^11–14^. Several reasons have been hypothesized for the failure of such trials, including multi-organ failure due to the underlying disease^15^, poor performance of current surfactant preparations relative to natural lung surfactant^16,17^, inactivation of lung surfactant by serum proteins^11,18^, delayed administration of the surfactant therapy^17^ and inadequate delivery to the alveolar regions of the lungs^19–21^.

Improving the delivery of exogenous surfactant to the alveolar regions has been deemed feasible by optimizing the volumetric dose of delivered surfactant (i.e. 2-4 ml/kg body weight compared to 1 or 8.5 ml/kg in the aforementioned studies), thus compensating for the amount of surfactant lost along the conducting airway walls, while avoiding adverse effects due to high doses^20^. Nevertheless, it remains uncertain whether adjusting solely for the surfactant doses would ultimately improve treatment outcomes. For example, significant heterogeneous surfactant distribution was observed in sheep^22,23^ and pig^24^ lungs following liquid instillations, despite using adequate doses (i.e. 4 ml/kg). Such outcomes may suggest that instillations of liquid boluses tend to leave some lung regions untreated while dangerously overflooding others. The leading posited reason underlying inhomogeneous distribution in large lungs compared to a relatively uniform distribution in smaller neonatal lungs follows from increased gravitational effects on liquid plug propagation that are proportional to the square of the characteristic length scale^20^ (e.g. airway diameter). In parallel to such efforts, alternative surfactant therapy strategies have included aerosol delivery of surfactant that has shown improved distribution compared to liquid instillations^22^. While inhalation therapy is currently pursued as a treatment option for infants with RDS^25,26^, its applicability for ARDS in adults has been largely hampered by low aerosol delivery efficiencies^6,27–29^.

Motivated by the ongoing lack of effective pharmaceutical interventions in ARDS^30^, we developed a novel surfactant delivery method (liquid foam therapy or LiFT) that involves foaming the surfactant prior to intratracheal administration in order to deliver it more homogenously throughout the lungs. First, we compared the distribution of a foamed versus liquid surfactant surrogate solution in *ex vivo* adult pig lungs. We hypothesized that foam would distribute more uniformly within large lungs compared to liquid due to its increased volume and lesser tendency to drain along the direction of gravity; a characteristic that follows from its high effective viscosity coupled with a low density^31^. Next, we tested the safety and efficacy of a foamed lung surfactant preparation in an *in vivo* rat model of ARDS induced by repeated whole lung lavage^32^. We posited that foamed lung surfactant has similar efficacy as liquid instillations in a well-established small animal ARDS model, where liquid instillations are known to be effective. Our efforts represent a significant first validation step of LiFT towards large animal studies and clinical trials.

## Results and discussion

### *Ex vivo* assays in excised adult pig lungs

To compare the distribution of foamed and liquid lung surfactant following intratracheal administration into large lungs, we first developed an *ex vivo* experimental model in ventilated excised pig lungs (see Fig. 1a and Methods). In these experiments, an affordable dyed lung surfactant surrogate (DLSS) solution (normal saline with 14% Tween 80 and 1% green dye) was used as a substitute for costly animal-derived lung surfactants sold commercially (~$100/ml). This solution is easy to visualize and has a kinematic viscosity similar to that of Infasurf; a commercial surfactant preparation used in our *in vivo* experiments (see Methods). DLSS was readily foamed using a hand mixer and the resulting foam had bubble sizes in the range of ~50 to ~800 µm with a half-drainage time of ~2.5 min (see Methods and Supplementary Material Fig. S3). The DLSS solution was administered to the lungs either as liquid (1 ml/kg, n=5) or foam (1 ml/kg before foaming, n=5), and the lung lobes were separated, sliced and imaged (see Methods). Since surfactant distribution depends heavily on volume doses and administration procedures, comparing foam and liquid administration is intrinsically challenging. Thus, we chose to follow closely protocols of recent clinical trials^11,13^ and explore whether surfactant distribution in such trials could be improved by foaming the surfactant prior to administration.

**Figure 1.**
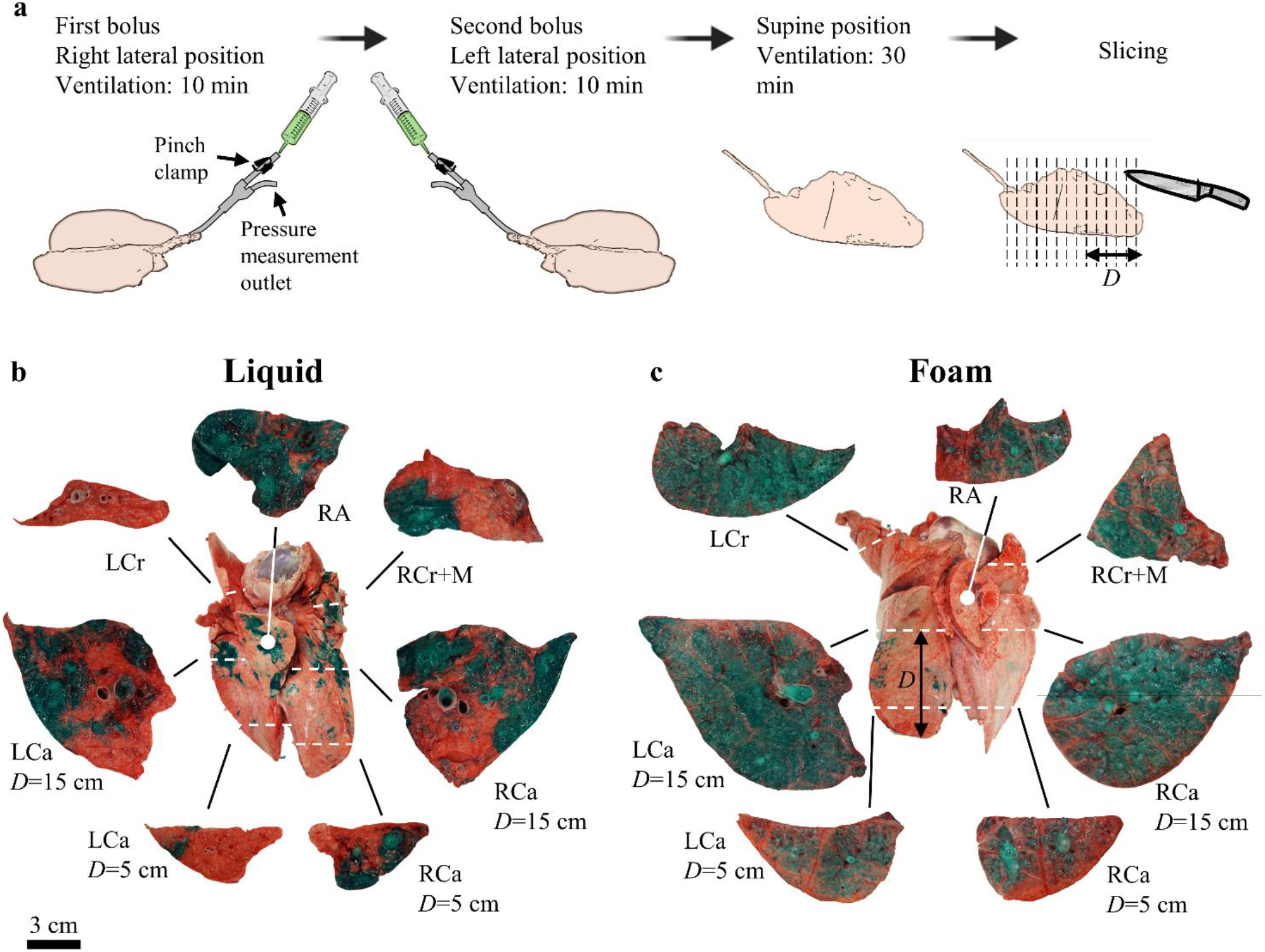
*Ex vivo* experiments. (**a**) Experimental protocol: healthy lungs from a ~90 kg pig are fully recruited, placed in a right lateral position and mechanically ventilated (7 ml/kg, 12 bpm, PEEP = 5 cmH_2_O, I:E ratio=1:2). Ventilation is paused at end-expiration and a DLSS solution (14% w/w Tween 80, 1% w/w green histological dye in normal saline) is administered to the lungs as either liquid (0.5 ml/kg) or foam (0.5 ml/kg before foaming, 3.1 ml/kg after foaming using a hand mixer), followed by a 3 ml/kg air bolus. The lungs are then further ventilated for 10 min. Next, a second 0.5 ml/kg bolus is similarly administered at a left lateral position followed by a 3 ml/kg air bolus and 10 min of ventilation. Then, the lungs are ventilated for 30 min in the supine position. Finally, the lung lobes are separated and cut along the transverse direction into 2.5 cm thick slices (see Methods). (**b,c**) Visualization of *ex vivo* distribution: example images of lungs and corresponding lung slices following administration of (**b**) liquid and (**c**) foamed DLSS. For each lung, the ventral side of the full lung is shown alongside example lung slices from the left cranial (LCr), left caudal (LCa), right caudal (RCa), right cranial+right middle (RCr+M) and right accessory (RA) lobes. White dashed lines indicate the approximate location of the shown slices within the lung. *D* corresponds to the measured distance from the caudal edge of the caudal lobe to the imaged face of the slice. Note that the scale bar (3 cm) corresponds to the lung slices and that the full lungs (shown in the center of **a** and **b**) are mirrored and not shown to scale.

### Dye distribution imaging

Figure 1 shows examples of lungs and lung slices following administration of liquid (Fig. 1b) and foamed (Fig. 1c) DLSS solution. Liquid DLSS exhibited a patchy distribution where some lung regions received high quantities of dye while other regions, as large as a few centimeters across, remained essentially void of dye. Moreover, the entire left cranial lobe (LCr) received very little dye in all (n=5) lungs treated with liquid DLSS (see Fig. 1b and discussion below). These results agree well with previous *in vivo* experiments that showed patchy pulmonary surfactant distribution^24^. In contrast, foamed DLSS reached all lung lobes and was more homogenously distributed throughout the tissue, typically leaving only millimeter-sized gaps between dyed areas. In both foam and liquid treatments, we observed reduced dye deposition in the caudal (and more distal) side of the caudal lobes compared to the cranial (more proximal) side.

### Quantitative analysis of *ex vivo* dye distribution

Dye distribution patterns for foamed and liquid DLSS were compared quantitatively by calculating the percentage of dyed area in the lung-slice images (dyed fraction) using image analysis (see Fig. S1). The average dyed fraction for the whole lung (Fig. 2a) was 53±3% for the foam group and 39±2% for the liquid group (95% CI for the difference: 6 to 23%, p=0.005). A higher dyed fraction for the foam group was also observed in the left caudal (p=0.01), left cranial (p=0.01), and right cranial (p=0.02) lobes. Conversely, the dyed fraction in the right accessory lobe was higher for the liquid group. However, the high variability of liquid DLSS delivery in this small lobe reflects an “all or nothing” scenario and precludes statistical inference (p=0.26). Interestingly, the average dyed fraction was similar in the right caudal lobe for the foam (48±6%) and liquid (52±7%) groups (95% CI for the difference: −18 to 26%, p=0.7). This may reflect a redistribution of DLSS present in the right caudal lobe during the 10 min of ventilation in the left lateral position. The inter-lobar standard deviation was 29±4% in the liquid group and 16 ±4% in the foam group (95% CI for the difference: 0.5 to 25%, p=0.04), indicating a more uniform inter-lobar distribution in the foam group.

**Figure 2.**
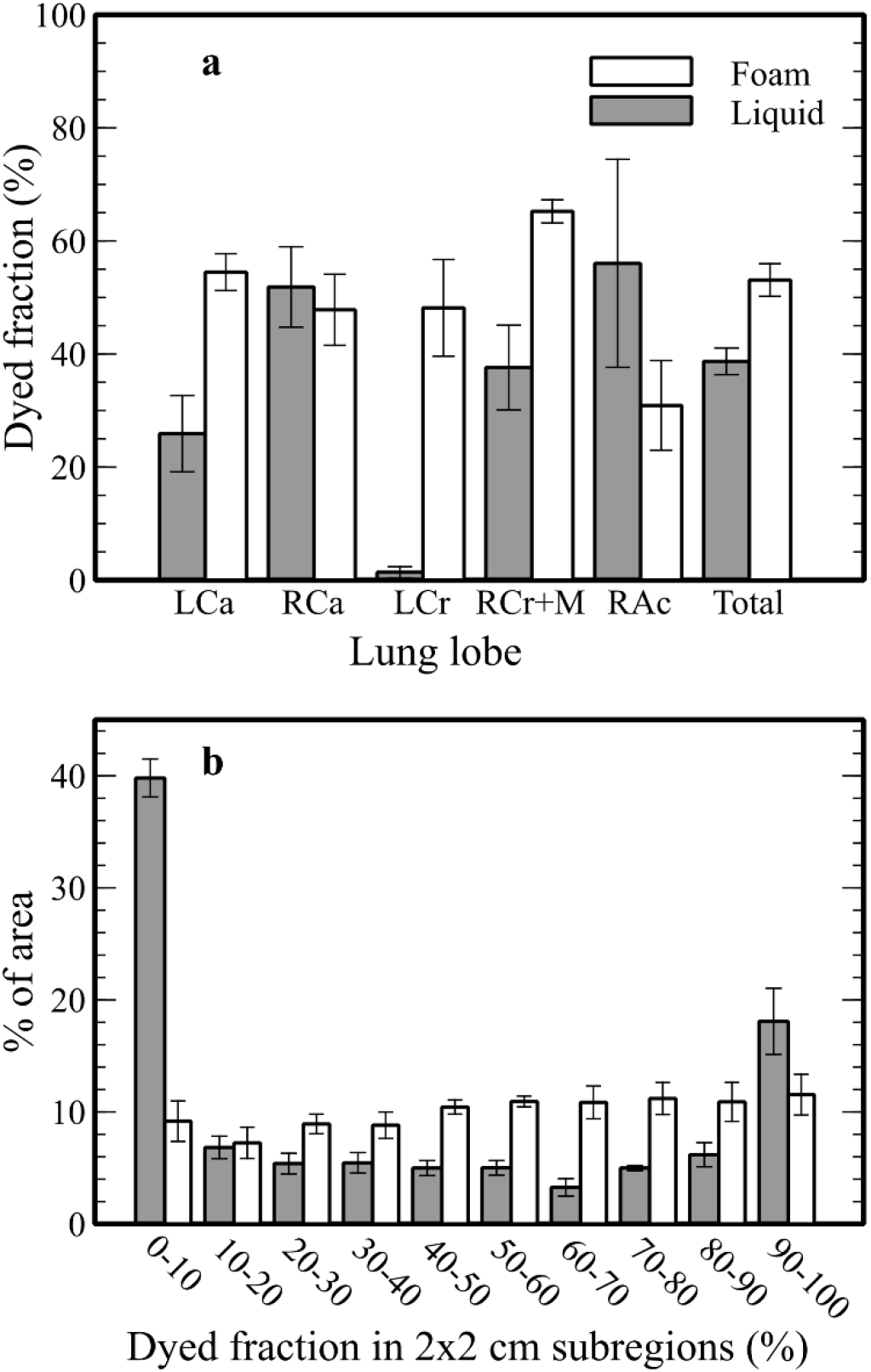
Quantification of *ex vivo* dye distribution. (**a**) Mean (±SE) dyed fraction for individual lobes and the whole lung following *ex vivo* administration of foamed (1 ml/kg before foaming, n=5) and liquid (1 ml/kg, n=5) DLSS. The dyed fraction is defined as the percentage of dyed surface area in the lung slices (e.g. as shown in Fig. 1b,c). The total dyed fraction was higher by 14.4% in the foam group (95% CI = 6-23%, p=0.005). This indicates a more uniform dye distribution in the foam group compared to the liquid group since the DLSS dose was fixed. The inter-lobar standard deviation of the dyed fraction was higher by 12.8% in the liquid group (95% CI = 0.5-25.2%, p=0.04) suggesting more equal partitioning of DLSS between lung lobes for the foam group. (**b**) Normalized dye distribution in sub-regions of the lung slices created by digitally superimposing a 2×2 cm grid on the original slice images (see Methods and SM). Values represent the mean (±SE) percentage of sub-regions across 10% distribution intervals, weighted by the area of the sub-region. The percentage of lung slice area in sub-regions with a dyed fraction > 10% (i.e. sum of the 9 right columns shown in the histogram) was 91±2% for the foam group and only 60±2% for the liquid group (95% CI for the difference: 25 to 36%, p<0.001).

While the higher total dyed fraction in the foam group indicates a more uniform overall distribution, this analysis alone does not fully capture the qualitative differences in dye patterns observed in lung slice images (Fig. 1b,c). To quantitatively characterize these differences, we digitally divided the lung-slice images into smaller sub-regions (see Methods) and calculated the dyed fraction for each sub-region (Fig. S1). We then constructed a normalized histogram showing the number of sub-regions for each 10^th^ percentile of dyed fraction, weighted by the area of the sub-region (Fig. 2b). In the liquid group, 40±2% of the lung slice area was found in sub-regions with a dyed fraction < 10% reflecting the large untreated areas observed qualitatively. In contrast, only 9±2% of the area resided in sub-regions with a dyed fraction < 10% for the foam group indicating that untreated regions therein are typically small. This observation may be critical for surfactant treatment in adults since surfactant is known to spread promptly across small lung regions^33^ but is less likely to spontaneously distribute across larger regions. It is thus possible that the small untreated gaps in the foam treatment would be bridged by the spontaneous spreading kinetics of surfactant, while the larger gaps in the liquid treatment would remain untreated. We underline that the percentage of area in sub-regions with a dyed fraction > 10% (i.e. sum of the nine right columns of Fig. 2b) was 91±2% for the foam group and only 60±2% for the liquid group (95% CI for the difference: 25 to 36%, p<0.001); a difference that may highly influence the efficacy of surfactant treatment in adults. Note that lobes receiving a small or no amount of DLSS were smaller in size than those containing high amounts of DLSS (e.g. left cranial lobes in Fig. 1b,c). Since this bias leads to an underestimation of the contribution of large untreated regions in the analysis, our experiments are likely to overestimate the total dyed fraction in the liquid group.

To quantitatively analyze the lower dyed fraction observed for more distal slices in the caudal lobes (Fig. 1b,c), we compared the total dyed fraction for each slice at different distances *D* (Fig. 1) from the caudal edge of these lobes to the imaged face of the lung-slices (Fig. 3). Notably, the average dyed fraction for the range *D*=12.5 to 17.5 cm (proximal region) was higher than the average dyed fraction in the range *D*=2.5 to 10 cm (distal region) for the foam group in the left caudal lobe (difference: 23±4%, p=0.004), as well as for the liquid group in the left caudal lobe (difference: 23±8% p=0.049), and the foam group in the right caudal lobe (difference: 24±6%, p=0.02) but not for the liquid group in the right caudal lobe (difference: −0.5±12%, p=0.97). The difference between dyed fraction in proximal vs distal regions did not differ significantly between the treatments (p=0.9 and p=0.1 for the left and right caudal lobes, respectively). Such decreased levels of surfactant delivery to the distal regions of the caudal lobes were previously reported in *in vivo* experiments using natural bovine surfactant^24^. In the liquid group, this decrease is most likely a result of surfactant loss along the airway walls and alveolar filling in more proximal lung regions^20^. In the surfactant group, the increased effective viscosity of the foam may have resulted in the obstruction of distal airways (see below) and a preferred distribution to more proximal regions that are reached through shorter paths. Such sub-optimal foam distribution in distal lung regions may potentially be improved by increasing the dose above 1 ml/kg or including a post-treatment recruitment maneuver.

**Figure 3.**
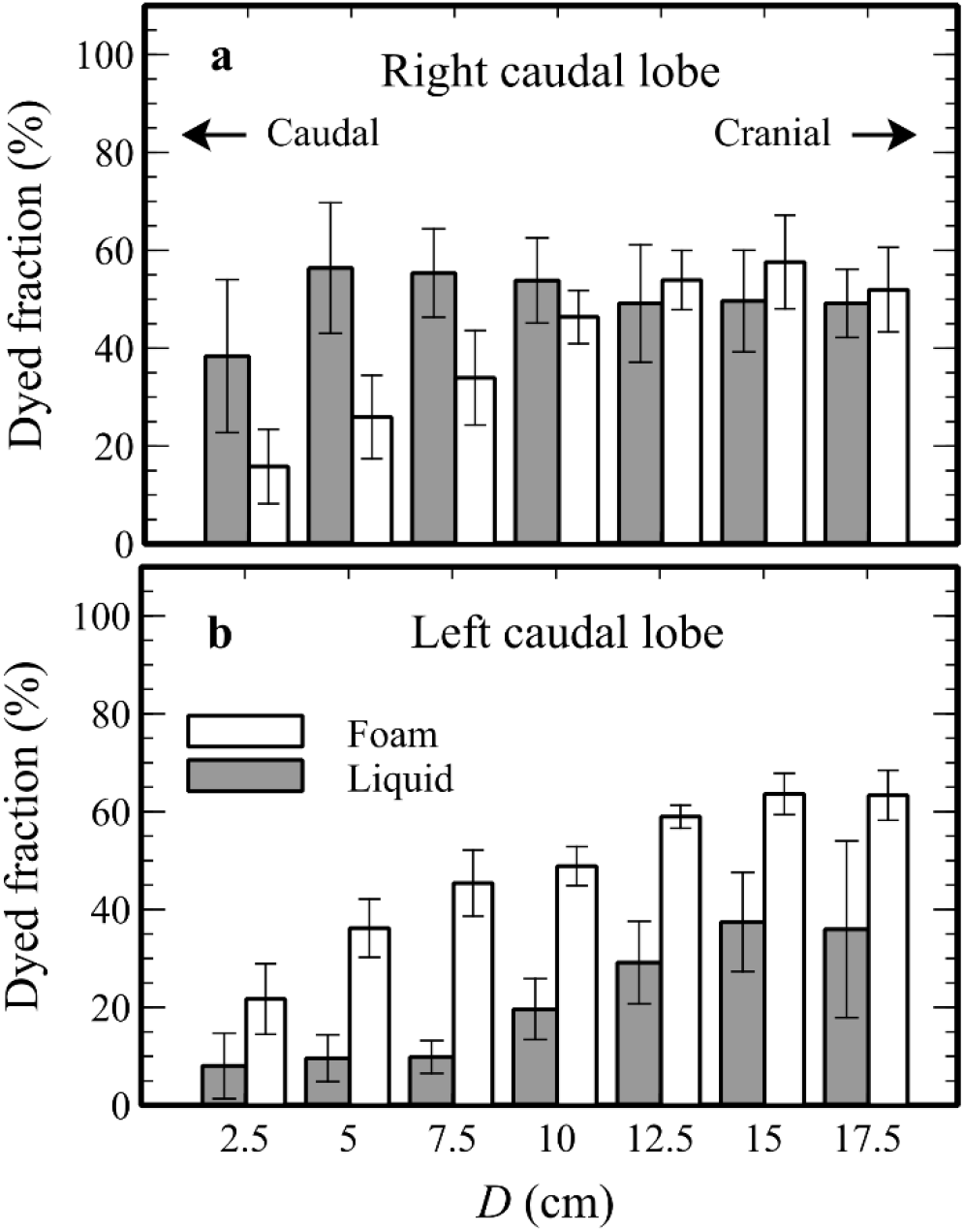
*Ex vivo* caudal-cranial distribution in the caudal lobes. Mean (±SE) dyed fraction within lung slices for the (**a**) right and (**b**) left caudal lobes following administration of liquid or foamed DLSS. Values are presented as a function of the distance *D* measured from the caudal edge to the imaged face of the slice (see Fig. 1a,c). The average dyed fraction for *D*=12.5 to 17.5 cm was higher than the average dyed fraction for *D*=2.5 to 10 cm for the foam group in the left caudal lobe (p=0.004), for the liquid group in the left caudal lobe (p=0.049), and for the foam group in the right caudal lobe (p=0.02).

### *Ex vivo* pressure measurements

Mean and peak pressure values before, during and after liquid and foam administration in the *ex vivo* experiments are summarized in Table 1. The peak inspiratory pressure (PIP), measured as the mean of peak pressure values in 30 sec, was similar before the treatment in the liquid group (10.1±0.2 cm H_2_O) and the foam group (11.0±1.1 cmH_2_O). Five min after the first bolus, the PIP value in the foam group was 11.1 cmH_2_O higher, on average, in the foam group compared to the liquid group (PIP_foam_=23.5±0.4 cmH_2_O, PIP_liquid_=12.4±1.1 cm H_2_O, p<0.001). However, the mean interval in PIP value between the groups reduced to 8.8 cmH_2_O 5 min after the second bolus and decreased further to 5.2 cmH_2_O 40 min after the second bolus. This reduction in PIP interval is a result of an increase in PIP for the liquid group following the second bolus compared to its value after the first bolus (p=0.046, paired t-test) and a reduction in PIP in the foam group between 5 and 40 min after the second bolus (p=0.005, paired t-test). The reduced PIP in the final ventilation period may be attributed to a gradual displacement of the foam towards distal generations, along with the opening of foam plugs due to breakage (i.e. rupture of foam bubbles) and drainage.

**Table 1.** Measurements of *ex vivo* pressures and treatment duration.

|  | Foam | liquid |
| --- | --- | --- |
| Total treatment duration first bolus (sec) | 258±11 | 75.5±6** |
| Total treatment duration second bolus (sec) | 293±16 | 60±5** |
| PIP before treatment (cm H<sub>2</sub>O) | 10.1±0.2 | 11.0±1.1 |
| PIP 5 min after first bolus (cm H<sub>2</sub>O) | 23.5±0.4 | 12.4±1.1** |
| PIP 5 min after second bolus (cm H<sub>2</sub>O) | 26.6±0.8 | 17.8±1.2** |
| PIP 40 min after second bolus (cm H<sub>2</sub>O) | 23.3±1.1 | 18.1±1.2* |
| Average P during first bolus (cm H<sub>2</sub>O)† | 26.0±0.7 | 4.3±0.9** |
| Average P during second bolus (cm H<sub>2</sub>O)† | 28.0±0.5 | 4.9±1.0** |
| Peak P during first bolus (cm H<sub>2</sub>O) | 47.5±1.6 | 7.4±1.0** |
| Peak P during second bolus (cm H<sub>2</sub>O) | 51.6±2.6 | 8.6±1.2** |

\*\*p<0.001
\*p=0.047
†Excluding time between the two injections for foam group

Time-averaged and peak pressures measured outside the endotracheal tube during foam injection were higher in the foam group compared to the liquid group in both boluses (p<0.001, Table 1). However, for all foam administration sessions, the time-averaged pressure was lower than 30 cmH_2_O, and the peak momentary pressure never exceeded 60 cmH_2_O; such values are not expected to result in significant barotrauma to the lungs. For comparison, peak pressures are regularly raised to 40-60 cmH_2_O during recruitment maneuvers in ARDS patients^34^. Furthermore, the measured pressure values represent an overestimation of pressure inside the lung since the high effective viscosity of the foam increases pressure drop across the endotracheal tube and the pressure was measured outside the endotracheal tube. Nevertheless, an extended period of time was required to maintain low pressures while injecting foam (~5 min) compared to liquid (~1 min, Table 1). This could indicate a need to balance potentially harmful pressures during foam injection with the risk of extending the time period when a patient is not being ventilated. The increased PIP following foam injection is likely a result of the relatively high pressures required for propagating foam plugs in small airways due to the foam’s high effective viscosity^31,35^. Such foam plugs may also entirely obstruct lung airways, thus reducing the volume of the lung available for ventilation. This is in line with reduced foam distribution to the distal part of the caudal lobes (Fig. 3). Nevertheless, these foam plugs seem to open up over time as witnessed in the gradual reduction in PIP within 40 min post-administration (Table 1).

It should be noted that our *ex vivo* pressure measurements do not directly reflect anticipated outcomes of LiFT in *in vivo* models or ARDS patients. First, a decrease, rather than an increase in lung pressure is expected after the treatment due to the effectivity of exogenous lung surfactant in lowering surface tension and reducing inflammation (see *in vivo* experiments below). Additional factors may also affect lung pressures *in vivo* that are not reflected in our *ex vivo* model, e.g. alveolar fluid clearance^36^ and autonomous breathing when a muscle relaxant is not used, which could pull the foam deeper into the lung. Finally, foam characteristics (i.e. effective viscosity, bubble sizes and foam stability) may differ between DLSS and a pulmonary surfactant (see Methods and Fig. S3). Nevertheless, our *ex vivo* measurements provide a strong initial indication that LiFT can dramatically improve the homogeneity of surfactant distribution in large lungs while maintaining a safe range of pressures during administration and subsequent ventilation.

### *In vivo* surfactant depletion model

We assessed the *in vivo* efficacy of foamed calf lung extracted surfactant (Infasurf; ONY, Inc., NY) using a well-established model of ARDS in rats^32^. Seventeen rats were anesthetized using standard procedures and lung injury was induced by repeated whole lung lavage with warm saline followed by 30 min of ventilation to ensure model stability. Animals were then randomly treated with foamed Infasurf (3 ml/kg before foaming, 12 ml/kg after foaming, n=6) followed by 1 ml of air, liquid Infasurf (3 ml/kg, n=6) followed by an air bolus, or sham treatment with air as control (n=5). The air bolus following liquid treatment and the volume of the sham air-treatment were adjusted to equate the total volume in all three groups. Infasurf foaming was achieved using a modified Tessari double-syringe technique^37,38^, originally developed for foam sclerotherapy (see Methods). The resulting foam had a smaller bubble size range of ~5 to ~1000 μm (Fig. S3) and an increased half-drainage time of ~30 min compared with DLSS foam.

### Arterial PO2 (PaO_2_) measurements

As a result of the lavage-induced lung injury, the mean PaO_2_ value for the 17 rats included in this study reduced from a baseline of 442±6 mmHg to 69±3 mmHg 5 minutes after the last lavage, indicating severe hypoxemia (Fig. 4a). 30 min after the last lavage PaO_2_ levels remained stable at a mean value of 70±4 mmHg. Mean PaO_2_ values increased dramatically within 15 min after treatment to >320 mmHg both in the foam and liquid treatments. These values remained stable until the end of the experiment (i.e. 90 min post-treatment). In the sham treatment, PaO_2_ values in two rats did not surpass 100 mmHg after treatment, while for the other 3 rats a gradual increase in PaO_2_ was observed; this latter improvement may be due to alveolar recruitment induced by the air bolus^39^.

**Figure 4.**
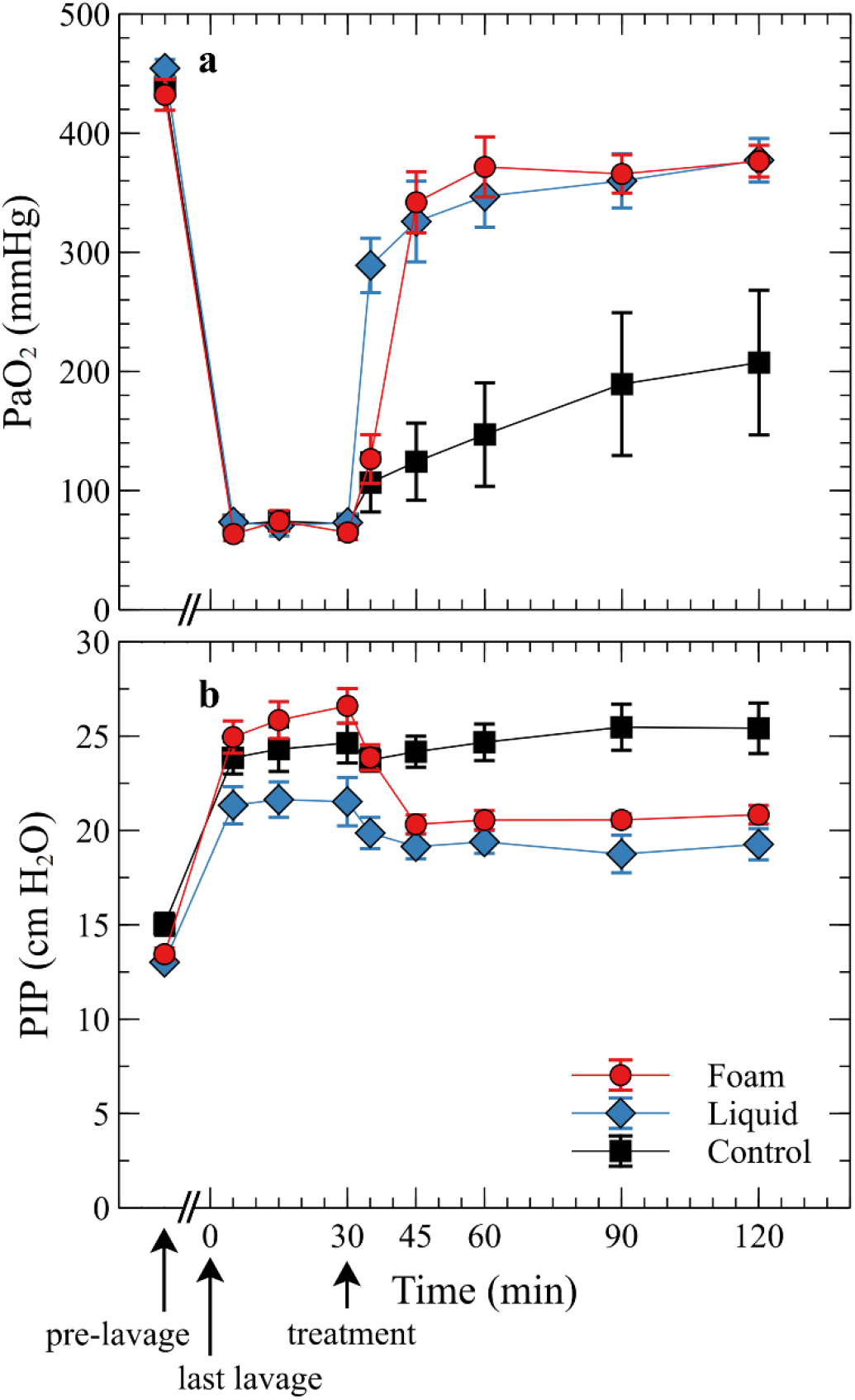
*In vivo* results in rats. (**a**) Mean (±SE) PaO_2_ values for animals receiving liquid Infasurf (3 ml/kg, n=6), foamed Infasurf (3 ml/kg foamed to 12 ml/kg, n=6), or sham treatment with air as control (n=5). Excess area under the PaO_2_ curve spanning treatment to 90 min post-treatment (AUC_PaO2_) was higher in the foam group (p<0.001) and liquid group (p<0.001) versus the sham group; note that the foam group did not differ from the liquid group (95 percent confidence for the difference: −7186 to 8046 mmHg×min or −29.8 to 33.4% of the mean AUC_PaO_2__ for the liquid group, p=0.99). The mean PaO_2_ value 5 min post-treatment was higher in the liquid group compared to both the foam group (p<0.001) and the control group (p<0.001), indicating a slower response with the foam treatment. **(b)** Mean (±SE) PIP values for the liquid, foam and control groups. Area between the PIP curve and a horizontal line representing the pre-treatment value spanning treatment to 90 min post-treatment (AUC_PIP_) was higher in the foam group compared to the control group (p<0.001).

Following the methodology used in clinical trials^9,11^, we assessed the overall effect of the different treatments on PaO_2_ by calculating the area between the graph of measured values and a horizontal line representing the pre-treatment value measured prior to treatment (AUC_PaO_2__). The mean difference in AUC_PaO_2__ value between the foam group and the control group was 16,399 mmHg×min or ~202% of the mean AUC_PaO_2__ value for the control group (95 percent confidence interval 8,412 to 24,387 mmHg×min or 103 to 300% of the mean AUC_PaO_2__ value for the control group, p<0.001). Similarly, the mean difference in AUC values between the liquid and control groups was 15,969 mmHg×min or 196% of the mean AUC_PaO_2__ value for the control group (95 percent confidence interval 7,982 to 23,957 mmHg×min or 98 to 294% of the mean AUC_PaO_2__ value for the control group, p<0.001). The mean difference in AUC values between the foam and liquid groups was 430 mmHg×min or 1.8% of the mean AUC for the liquid group (95 percent confidence −7186 to 8046 mmHg×min or −29.8 to 33.4% of the mean AUC_PaO_2__ for the liquid group, p=0.99). Thus, both liquid and foam treatments were effective in increasing PaO_2_ levels, and this effect was very similar for the two treatments. 5 min post-treatment administration the mean PaO_2_ value for the liquid group (PaO_2_=289±23 mmHg) was higher compared to the foam group (PaO_2_=126±20 mmHg, p<0.001) and compared to the control group (PaO_2_=107±25 mmHg p<0.001), while the mean PaO_2_ value for the foam group was similar to that of the control group (p=0.8). This indicates that improvement in PaO_2_ was somewhat slower in the foam group compared to the liquid group. Further results for dynamic compliance, arterial PCO2 and pH measurements are presented in Fig. S2.

### *In vivo* pressure measurements

The mean PIP value for all rats increased from a baseline value of 13.8±0.3 cm H_2_O to 23.5±0.6 cm H_2_O 5 min after the last lavage and remained stable until the treatment (Fig. 4b). Importantly, pre-treatment PIP values differed between the groups (26.6±0.9, 21.5±1.3 and 24.6±1.1 cm H_2_O for the foam, liquid and control groups, respectively). However, these differences are fortuitous since randomization took place after the pre-treatment measurements. 15 min after the treatment, the PIP value in the foam group reduced to 20.3±0.5 cm H_2_O and remained stable until the end of the experiment. This is in contrast to the rise in PIP observed *ex vivo* and suggests that the foamed Infasurf helped in opening collapsed regions of the lung and/or reducing surface tension. A moderate reduction in PIP (to 19.9±0.8) was also observed in the liquid group, but PIP did not change appreciably in the control group. The area between the PIP curve and a horizontal line representing the pre-treatment value from treatment to 90 min after treatment (AUC_PIP_) was higher in the liquid (AUC_PIP_=217±75 cmH_2_O×min, p=0.075) and foam (AUC_PIP_=487±72 cmH_2_O×min p<0.001) groups compared to the control group (AUC_PIP_=46±46 cmH_2_O×min), and higher in the foam group compared to the liquid group (p=0.017). To correct for potential bias induced by the differences in pre-treatment values we also compared AUC_PIP_ between the different groups using analysis of covariance with pretreatment PIP as a covariate; we find p<0.001 for liquid vs control, p<0.001 for foam vs control, and p=0.7 for foam vs liquid. Overall, these results are in line with our PaO_2_ measurements and show that the foam and liquid treatments can reduce PIP *in vivo*. Furthermore, the decrease in PIP following foamed Infasurf administration suggests that the rise in PIP observed *ex vivo* may not be predictive of potential barotrauma risks *in vivo*. Further results for dynamic lung compliance are presented in Fig. S2.

### Outlook

Over the past decades, liquid instillations have become the mainstay for exogenous surfactant delivery in premature infants. Yet, recent studies indicate major disadvantages of liquid instillations when applied to adult ARDS patients, such as a patchy, non-uniform distribution. Here, we showed that LiFT results in a more uniform pulmonary distribution than liquid instillations in large *ex vivo* lungs when using a surrogate lung surfactant solution. In addition, foamed Infasurf was able to increase PaO_2_ and reduce PIP in an *in vivo* rat model of ARDS. These findings suggest that LiFT may greatly improve the efficacy of exogenous lung surfactant therapy in treating ARDS patients while being safe, thus advocating further investigation in large animal models towards new and more efficient surfactant treatments for ARDS. Furthermore, using LiFT as a drug carrier for topical delivery of drugs and cells to the lungs may hold tremendous potential for a wider range of pulmonary therapies^40,41^.

## Methods

### Solution preparation and characterization

Dyed lung-surfactant surrogate (DLSS) solution was prepared by mixing a green histological marking dye (Wak-Chemie Medical, Germany) and Tween 80 solution (14% w/w in normal saline) at a 1:100 mass ratio. To avoid precipitation of the dye, the DLSS solution was mixed using a magnetic stirrer right before use. The kinematic viscosity of the solution before adding dye was 3.0 cSt at 20°C, as measured using a size 50 Cannon Fenske viscometer (Cannon Instrument Co., PA, and Cole Parmer Instrument Co., IL, average of two independent measurements). This value corresponds roughly to the kinematic viscosity of Infasurf (2.5 cSt at 23°C to 3.9 cSt at 37°C)^42^.

### *Ex vivo* lung preparation

Ten pairs of lungs from ~90 kg adult pigs were purchased from a local provider (Lahav CRO, Israel) and kept at 4°C for up to two days and experiments were conducted at room temperature (22-25°C). To minimize potential damage to the lungs, the heart was not dissected from the lungs in the harvesting process and remained attached to them during the experiments. The lungs were ventilated using a PLV-100 ventilator (Respironics, USA) equipped with a single limb breathing circuit with an adjustable PEEP valve. This breathing circuit was connected to the lung via a silicone tube fitted with a ratchet pinch clamp, a three-way connector, and an 8.5 mm inner diameter endotracheal tube that was positioned at a 20-degree angle to the horizontal plane (Fig. 1a). Air pressure was monitored throughout the experiment via one outlet of the three-way connector using an in-house built pressure measurement system composed of a pressure transducer (AS-0023, Biometrix, USA), a voltage amplifier (ETH-256, iWorx, USA) and a data acquisition board (Arduino Uno, Arduino, Italy). This pressure measurement system was calibrated using a water column before each experiment. Full recruitment of the lungs was then established by setting a respiratory rate of 12 breaths per minute (bpm), with an inspiratory:expiratory time ratio (I:E ratio) of 1:2 and slowly (within ~20 min) increasing the tidal volume up to ~1.5 L and the PEEP up to ~20 cm H_2_O while alternately placing the lungs in the supine or prone position. Air-pressure did not exceed 60 cm H_2_O during the recruitment procedure. Subsequently, the PEEP and tidal volume were set to ~10 cm H_2_O and ~1L, respectively, and small superficial holes or rips in the lungs were clamped using hinge clips or tied with small cable ties. To mimic the *in vivo* orientation of the lung at a right lateral position, the lungs were positioned with the right lung facing down and the heart was propped on an adjustable lift stage and tied in place. The PEEP was then reduced to 5 cm H_2_O, the tidal volume was reduced to 0.64L (7 ml/kg), and the respiratory rate and I:E ratio remained unchanged. These settings were kept constant throughout the experiment. Note that the positioning of the lungs in a right lateral position was performed while maintaining higher tidal volume and PEEP than in subsequent steps to prevent de-recruitment of the lungs during manual handling.

### *Ex vivo* foam and liquid administration

Following the setup, the lungs were randomly assigned to one of two treatment groups: Liquid DLSS (1 ml/kg, n=5) or foamed DLSS (1 ml/kg before foaming, ~6.2 ml/kg after foaming, n=5). Both treatments were divided into two aliquots, such that the first half of the dose was administered in a right lateral position, while the second half was given in a left lateral position. In the foam group, 51 gr of DLSS were foamed inside a graded plastic beaker (inner diameter 5 inches) using a hand mixer at maximum power for 80 sec at room temperature (22-25°C) and then repeatedly foamed for an additional 10 sec (typically 1-3 times) until the volume of the foam reached ~325 ml. Before DLSS administration, the ventilation was stopped at end-expiration by closing the ratchet pinch clamp, and the ventilation circuit was disconnected from the silicone tube. Next, a 150 ml syringe was filled with 150 ml of foam by manually withdrawing foam from the beaker into the syringe, the tip of the syringe was inserted into the silicone tube, the ratchet pinch clamp was opened and the foam was injected into the lungs. An additional foam bolus of 150 ml was similarly administered immediately after the first one followed by two air boluses of 135 ml each that served to purge the foam from the tubing and conducting airways and propel it deeper into the lungs. An *in vitro* calibration performed before the experiments showed that this procedure produced a foam with a volume of ~280 ml at the distal end of the endotracheal tube, as measured using a measuring tube. This corresponds to a dose of ~3.1 ml of foam per kg body weight, and a total weight of 46 gr corresponding to a dose of ~0.5 ml of the original solution per kg body weight. To minimize the chance of barotrauma, the foam injection rate was adjusted to maintain a pressure of ~30 cm H_2_O. When injection was too slow, air pressure was allowed to rise up to ~40 cm H_2_O, with occasional peaks at ~50 cm H_2_O (see Results Table 1 for measured pressure values during foam and liquid administration).

Following the treatment, the lungs were reconnected to the ventilator and ventilated for 10 min at the right lateral position. Next, the lungs were turned to the left lateral position, and the administration procedure was repeated by applying an additional ~3.1 ml of foamed DLSS per kg (0.5 ml/kg before foaming) followed by two air boluses (135 ml each) and 10 min of ventilation in the left lateral position. Finally, the lungs were ventilated for 30 min in the supine position.

In the liquid group, 46 ml (~0.5 ml/kg) of liquid DLSS were injected into the lungs at the right lateral position followed by two air boluses of 135 ml each and 10 min of mechanical ventilation. This procedure was repeated in the left lateral position and was followed by 30 min of ventilation in the supine position.

### *Ex vivo* dye imaging and image analysis

Following foam or liquid administration, the lung lobes were separated using a sharp knife, and each lobe was sliced along the transverse plane at ~2.5 cm intervals. Since separating the right middle lobe from the right cranial lobe was difficult to perform accurately, these two lobes were not separated before slicing and were analyzed together as one unit. In addition, the accessory right lobe was sliced using a single cut orthogonal to the direction of the bronchus leading to this lobe. Note that the directions and orientations reported here relate to the geometry that the lungs assumed in this *ex vivo* experiment (see Fig. 1a) and that this geometry may differ from their shape *in vivo*. The slices were immediately imaged using a digital camera (Sony alpha A6000) with constant lighting and exposure settings, and a ruler was included in the frame as a reference for image scaling.

The pipeline for image analysis is depicted in Fig. S1. Images were digitally rescaled to correct for differences in the distance between the camera lens and the slices. Then, large cartilage regions, the lumen of large airways and the background outside the lung piece were masked out. The masked images were used to produce two binary images for each lung slice. In the first image (i.e. the ‘mask’ image) a value of 1 was assigned to all pixels that were not masked out in the previous step and 0 was assigned otherwise. In the second image (the ‘dye’ image) pixels containing green dye were assigned a value of 1 and pixels without green dye were assigned a value of 0. This image was obtained by first applying a transformation of RGB color space into Lab color space^43^ and then manually selecting a threshold for the ‘a’ channel that results in the best segmentation of the green dye. This resulted in a reasonably good segmentation of green areas from red ones since in Lab color space the ‘a’ axis extends from green to red. The operator selecting this threshold value was blinded to the treatment group. Next, the two binary images were used to determine lobar and caudal-cranial dye distributions. The dyed fraction for each lung slice was determined by dividing the sum of pixel values in each dye image by the sum of pixel values in the corresponding mask image and multiplying by 100. The lung slices were then digitally divided into sub-sections by subdividing the mask and dye images into 2×2 cm macropixels (Fig. S1g). The resulting dyed fraction in each sub-section of the lung was calculated by dividing the sum of pixel values in each macropixel of the dye image by the sum of pixel values in the corresponding macropixel of the mask image and multiplying by 100. The area of the subsection was calculated by multiplying the sum of values in a macropixel of the mask image by a scale factor based on the size calibration. A histogram was then created representing the number of subsections for each 10^th^ percentile of the dyed fraction, weighted by the area of the subsection.

### Animal preparation

All procedures were approved by the ethical committee of the Technion-Israel Institute of Technology (permit no. IL-132-11-18). Adult male Wistar rats (370-550 gr) were first lightly anesthetized by isoflurane inhalation and then fully anesthetized by an intraperitoneal (i.p.) injection (75 mg/kg ketamine and 5 mg/kg xylazine in sterile normal saline). Next, the rats were placed supine on a warm plate and covered with a blanket to maintain normal body temperatures (36-38°C), the tail vein was cannulated (BD Neoflon, 24GA or 26GA, Becton Dickinson Infusion Therapy AB, Sweden), and anesthesia was maintained by continuous intravenous (i.v.) injection of Propofol (10 mg/ml, up to 5.4 ml/kg/h). When needed, additional injections of anesthetic drugs were given (preferably propofol 10 mg/ml, up to a 1 ml/kg bolus, i.v.; or a mix of 25 mg/kg ketamine and 1.3 mg/kg xylazine; or 1.3 mg/kg xylazine alone). When sedation was deep, the continuous injection of propofol was temporarily replaced by normal saline. Importantly, this liquid administration regime was calibrated to avoid large drops in blood pressure. Buprenorphine (0.03 mg/ml, 0.2 ml) was given subcutaneously for analgesia. Bupivacaine was applied to the skin at the throat area for additional local anesthesia. Subsequently, the carotid artery was exposed and cannulated (BD Neoflon, 26GA) to allow blood pressure measurement and arterial blood sampling (a small spatula with inverse adhesive tape was inserted under the carotid to anchor it during cannulation). To avoid blood clotting in the tubes, the cannula was first washed with a 0.2 ml bolus of a dilute heparin solution (9 U/ml in normal saline) and then continuously perfused with the same solution at 0.3 ml/h for the rest of the experiment. Subsequently, a tracheostomy was performed and the trachea was intubated with a 14-gauge blunt syringe needle. The rats were ventilated using a Harvard small animal ventilator (Harvard Bioscience, MA, USA) at a rate of 60 bpm, a tidal volume of 8 ml/kg, an I:E ratio of 1 and an oxygen fraction of 1 with the pressure limit set to 30 cm H_2_O. PEEP was maintained at 5 cm H_2_O throughout the experiment by immersing the exhaust tube in water. Atracurium besylate (2 mg/kg, i.v.) was administered to prevent spontaneous breathing. Additional injections of atracurium besylate (0.66 mg/kg i.v.) were given if spontaneous breathing resumed.

Following Lachmann *et al.*^44^, we measured the vital capacity of the rats’ lungs as follows: the rats were disconnected from the ventilation machine and air was slowly injected (within ~20 sec) into the lungs from a 20 ml syringe while monitoring the air pressure. Air injection was stopped when the pressure inside the lungs reached 35 cm H_2_O. The rats were then immediately reconnected to the ventilation machine and the volume of injected air (defined as the vital capacity) was recorded. Note that following this procedure, the average PIP for all rats reduced from 16.7±0.4 to 13.8±0.3 cm H_2_O. Subsequently, the lungs were ventilated for 10 min and baseline blood gas values were measured using a VetScan i-STAT 1 point-of-care analyzer (Abaxis, UK). Simultaneously, air pressure inside the lungs was recorded for 30 sec for assessing baseline PIP. The initial inclusion criterion was PaO_2_>370 mmHg. For lower baseline values, the respiratory rate was raised by 5 bpm and the tidal volume was raised by 1 ml/kg, rats were ventilated for an additional 10 min and baseline values were reassessed.

### Injury induction

Upon meeting the initial inclusion criterion, repeated whole lung lavage was performed to induce a severe lung injury. In each lavage, rats were disconnected from the ventilation machine and warm saline (37°C, volume=85% of the measured vital capacity) was injected into the lungs within ~9 seconds and withdrawn within ~11 sec. During saline injection, the pressure of injected saline was monitored and maintained below 35 cm H_2_O, using an analog blood pressure gauge, connected through a T connector. Immediately after each lavage, the rats were reconnected to the ventilation machine and the volume of retrieved saline was recorded. Model induction proceeded by the following steps: (i) 4 consecutive whole lung lavages were performed at 5 min intervals. (ii) Blood gas analysis was performed 5 minutes after the lavage. (iii) For PaO_2_>200 mmHg, an additional lavage was performed 10 min after the previous one. For 100 mmHg<PaO_2_<200 mmHg the lungs underwent lavage using half of the original lavage volume to avoid fatal lung damage. (iv) Steps ii and iii were repeated until the PaO_2_ value dropped below 100 mmHg. (v) The rats were further ventilated, and arterial blood gases and PIP were measured 15 and 30 min after the last lavage to ensure that PaO_2_ values are stable. (vi) Rats satisfying the experiment inclusion criterion of 45mmHg<PaO_2_<120 mmHg 30 min after the last lavage were included in further steps. If a PaO_2_ value higher than 120 mmHg was measured at 15 or 30 min after the last lavage, lavages resumed from step (iii).

### Treatment and monitoring

Rats were randomized to receive one of three treatments: (i) foamed Infasurf (foam group, 3 ml/kg before foaming, n=6) (ii) liquid Infasurf (liquid group, 3 ml/kg, n=6) or (iii) an air bolus (control group, n=5). In the foam group, Infasurf was foamed using a modified Tessari double-syringe technique^37,38^: first, a 5 ml syringe was filled with Infasurf (3 ml/kg + 0.2 ml to compensate for foam losses) and incubated for 5 min in a warm bath at 25°C. Next, the syringe was connected through a syringe filter (PES membrane, diameter 25 mm, pore size 5 μm) and a female-female Luer-lock connector to another 5 ml syringe. The volume of air in the second syringe was adjusted so that the air volume in the entire system (including 0.6 ml of air inside the filter and connector) was 3 times larger than the liquid volume. Finally, Infasurf was transferred 15-20 times between the two syringes, where the duration of each transfer was ~2 sec, resulting in dense and stable foam. The rats were then disconnected from the ventilation machine and the foam was slowly injected intratracheally within ~1 min. Based on prior calibration tests, this procedure resulted in the injection of 12 ml/kg of foam with a 1:3 liquid:air volume ratio. Immediately following foam injection, 1 ml of air was injected into the lungs from an adjacent syringe connected through a T connector to purge the foam from the tubes and conducting airways and push it deeper into the lungs. In the liquid group, a 2 ml syringe was filled with Infasurf (3 ml/kg + 0.1 ml to compensate for losses) and warmed for 5 min at 25°C in a warm bath. Infasurf was then injected into the lungs within ~10 sec followed by an air bolus (9 ml/kg + 1 ml, ~10 sec). In the control group, air (12 ml/kg + 1 ml) was injected within ~ 10 sec. In all three groups, pressure during the treatment was monitored using an analog pressure gauge and maintained below 35 cmH_2_O. Note that the overall volume injected was identical in all three groups to reduce confounding effects due to lung recruitment. Ventilation was resumed immediately after the treatment. In the foam group, ventilation immediately after the treatment was not optimal since the maximal pressure (set to 30 cm H_2_O) was reached. This resulted in a succession of short inhalation cycles that lasted ~1 min after which the ventilation resumed normally. Blood gases and PIP were measured at 5, 15, 30, 60 and 90 min after the beginning of treatment. Rats were then euthanized using an overdose of pentobarbital (200 mg/ml, 4ml/kg).

### Foam characterization

To compare the two types of foam used, we prepared foam samples using the same protocols as in the *ex vivo* and *in vivo* experiments (1 ml of Infasurf and 3 ml of air were used for the Infasurf foam). Bubble sizes were assessed by spreading a ~1 mm layer of foam on a glass slide with a syringe and imaging it in bright field mode at 2X and 10X magnification using a Nikon Eclipse Ti inverted microscope (Fig. S3).

Foam stability was characterized by measuring the foam drainage kinetics as follows: for the DLSS foam, a 5 ml syringe was filled with 5 ml of foam, the volume of the liquid fraction inside the syringe was measured by weighing the foam using a tared scale, the syringe was positioned vertically with the plunger facing downwards, and the time until half of the liquid drained to the bottom (i.e. half-drainage time) was measured. For the Infasurf foam, the syringe was placed vertically on its plunger immediately after the foaming process and the half-drainage time was measured as the time until 0.5 ml of liquid drained to the bottom.

### Statistical analysis

All data are presented as means±SE. Differences between 3 groups were determined using a one-way ANOVA and pair-wise P-values and confidence intervals were calculated using the Tukey-Kramer method. A paired two-sided t-test was used to compare values within groups. Differences between two groups were determined using an unpaired two-sided t-test and a Wilcoxon-Mann-Whitney test was used when assumptions for parametric testing were not met.

## Acknowledgments

This work was in part supported by the European Research Council (ERC) under the European Union’s Horizon 2020 research and innovation program (grant agreement No. 813228-LIFT); the Ministry of Science and Technology (grant Nos. 3-15649 and 3-16866); and the POLAK Fund for Applied Research, at the Technion. The authors wish to thank Rona Shofty, Amit Avrahami, Michal Schlesinger-Lau, Edith Cohen, Natalia Cohen, Tatyana Oriev, Osnat Nixon and Seffi Greenblat for their assistance.

## Notes

### Competing Interest Statement

R.F., Y. O., A.F., D.W. and J.S. are co-founders and shareholders of Neshima Medical Ltd.

## References

1. Umbrello, M., Formenti, P., Bolgiaghi, L. & Chiumello, D. Current Concepts of ARDS: A Narrative Review. Int. J. Mol. Sci. 18, 64 (2016).

2. Xu, Z. et al. Pathological findings of COVID-19 associated with acute respiratory distress syndrome. Lancet Respir. Med. 8, 420–422 (2020).

3. Gattinoni, L., Chiumello, D. & Rossi, S. COVID-19 pneumonia: ARDS or not? Crit. Care 24, (2020).

4. Lewis, J. F. & Jobe, A. H. Surfactant and the Adult Respiratory Distress Syndrome. Am. Rev. Respir. Dis. 147, 218–233 (1993).

5. Autilio, C. & Pérez-Gil, J. Understanding the principle biophysics concepts of pulmonary surfactant in health and disease. Arch. Dis. Child. Fetal Neonatal Ed. 104, F443–F451 (2019).

6. Raghavendran, K., Willson, D. & Notter, R. H. Surfactant Therapy for Acute Lung Injury and Acute Respiratory Distress Syndrome. Crit. Care Clin. 27, 525–559 (2011).

7. Jobe, A. H. Pulmonary Surfactant Therapy. N. Engl. J. Med. 328, 861–868 (1993).

8. Kwong, M. S., Egan, E. A., Notter, R. H. & Shapiro, D. L. Double-Blind Clinical Trial of Calf Lung Surfactant Extract for the Prevention of Hyaline Membrane Disease in Extremely Premature Infants. Pediatrics 76, (1985).

9. Spragg, R. G. et al. Effect of Recombinant Surfactant Protein C–Based Surfactant on the Acute Respiratory Distress Syndrome. N. Engl. J. Med. 351, 884–892 (2004).

10. Willson, D. F. et al. Effect of exogenous surfactant (calfactant) in pediatric acute lung injury: A randomized controlled trial. J. Am. Med. Assoc. 293, 470–476 (2005).

11. Spragg, R. G. et al. Recombinant surfactant protein C-based surfactant for patients with severe direct lung injury. Am. J. Respir. Crit. Care Med. 183, 1055–1061 (2011).

12. Willson, D. F. et al. Pediatric Calfactant in Acute Respiratory Distress Syndrome Trial. Pediatr. Crit. Care Med. 14, 657–665 (2013).

13. Willson, D. F., Truwit, J. D., Conaway, M. R., Traul, C. S. & Egan, E. E. The adult calfactant in acute respiratory distress syndrome trial. Chest 148, 356–364 (2015).

14. Kesecioglu, J. et al. Exogenous natural surfactant for treatment of acute lung injury and the acute respiratory distress syndrome. Am. J. Respir. Crit. Care Med. 180, 989–994 (2009).

15. Fuchs, L. et al. The Effect of ARDS on Survival: Do Patients Die From ARDS or With ARDS? J. Intensive Care Med. 34, 374–382 (2019).

16. Willson, D. F. & Notter, R. H. The future of exogenous surfactant therapy. Respir. Care 56, 1369–1386 (2011).

17. Rosenberg, Oleg A., Andrey E. Bautin, and A. A. S. Late start of surfactant therapy and surfactant drug composition as major causes of failure of phase III multi-center clinical trials of surfactant therapy in adults with ARDS. Int. J. Biomed. 8, 253–254 (2018).

18. López-Rodríguez, E., Ospina, O. L., Echaide, M., Taeusch, H. W. & Pérez-Gil, J. Exposure to polymers reverses inhibition of pulmonary surfactant by serum, meconium, or cholesterol in the captive bubble surfactometer. Biophys. J. 103, 1451–1459 (2012).

19. Kazemi, A. et al. Surfactant delivery in rat lungs: Comparing 3D geometrical simulation model with experimental instillation. PLOS Comput. Biol. 15, e1007408 (2019).

20. Filoche, M., Tai, C.-F. & Grotberg, J. B. Three-dimensional model of surfactant replacement therapy. PNAS 112, 9287–9292 (2015).

21. Grotberg, J. B., Filoche, M., Willson, D. F., Raghavendran, K. & Notter, R. H. Did reduced alveolar delivery of surfactant contribute to negative results in adults with acute respiratory distress syndrome? American Journal of Respiratory and Critical Care Medicine vol. 195 538–540 (2017).

22. Lewis, J. F. et al. Lung function and surfactant distribution in saline-lavaged sheep given instilled vs. nebulized surfactant. J. Appl. Physiol. 74, 1256–1264 (1993).

23. Lewis, J. F. et al. Evaluation of exogenous surfactant treatment strategies in an adult model of acute lung injury. J. Appl. Physiol. 80, 1156–1164 (1996).

24. Ruth Graham, M. et al. Quantitative computed tomography in porcine lung injury with variable versus conventional ventilation: Recruitment and surfactant replacement. Crit. Care Med. 39, 1721–1730 (2011).

25. Pillow, J. J. & Minocchieri, S. Innovation in Surfactant Therapy II: Surfactant Administration by Aerosolization. Neonatology 101, 337–344 (2012).

26. Cummings, J. J. et al. Aerosolized Calfactant for Newborns With Respiratory Distress: A Randomized Trial. Pediatrics 146, e20193967 (2020).

27. Anzueto, A. et al. Aerosolized Surfactant in Adults with Sepsis-Induced Acute Respiratory Distress Syndrome. N. Engl. J. Med. 334, 1417–1422 (1996).

28. Bianco, F. et al. From bench to bedside: In vitro and in vivo evaluation of a neonate-focused nebulized surfactant delivery strategy. Respir. Res. 20, 134 (2019).

29. Kim, H. C. & Won, Y. Y. Clinical, technological, and economic issues associated with developing new lung surfactant therapeutics. Biotechnol. Adv. 36, 1185–1193 (2018).

30. Matera, M. G., Rogliani, P., Bianco, A. & Cazzola, M. Pharmacological management of adult patients with acute respiratory distress syndrome. Expert Opin. Pharmacother. 1–15 (2020) doi:10.1080/14656566.2020.1801636.

31. Blauer, R., Mitchell, B., Regional, C. K.-S. C. & 1974, U. Determination of laminar, turbulent, and transitional foam flow losses in pipes. SPE Calif. Reg. Meet. Soc. Pet. Eng. (1974).

32. Czyzewski, A. M. et al. Effective in vivo treatment of acute lung injury with helical, amphipathic peptoid mimics of pulmonary surfactant proteins. Sci. Rep. 8, 1–9 (2018).

33. Davis, J. M., Russ, G. A., Metlay, L., Dickerson, B. & Greenspan, B. S. Short-term distribution kinetics of intratracheally administered exogenous lung surfactant. Pediatr. Res. 31, 445–450 (1992).

34. Bhattacharjee, S., Soni, K. D. & Maitra, S. Recruitment maneuver does not provide any mortality benefit over lung protective strategy ventilation in adult patients with acute respiratory distress syndrome: A meta-analysis and systematic review of the randomized controlled trials. J. Intensive Care 6, 35 (2018).

35. Hu, Y. et al. A microfluidic model to study fluid dynamics of mucus plug rupture in small lung airways. Biomicrofluidics 9, 044119 (2015).

36. Berthiaume, Y., Folkesson, H. G. & Matthay, M. A. Lung Edema Clearance: 20 Years of Progress Invited Review: Alveolar edema fluid clearance in the injured lung. J. Appl. Physiol. 93, 2207–2213 (2002).

37. Tessari, L., Cavezzi, A. & Frullini, A. Preliminary experience with a new sclerosing foam in the treatment of varicose veins. Dermatologic Surg. 27, 58–60 (2001).

38. Shirazi, A. R. & Goldman, M. The use of a 5-μm filter hub increases foam stability when using the double-syringe technique. Dermatologic Surg. 34, 91–92 (2008).

39. Frank, J. A. et al. Differential effects of sustained inflation recruitment maneuvers on alveolar epithelial and lung endothelial injury. Crit. Care Med. 33, 181–188 (2005).

40. Haitsma, J. J., Lachmann, U. & Lachmann, B. Exogenous surfactant as a drug delivery agent. Adv. Drug Deliv. Rev. 47, 197–207 (2001).

41. Hidalgo, A., Cruz, A. & Pérez-Gil, J. Barrier or carrier? Pulmonary surfactant and drug delivery. Eur. J. Pharm. Biopharm. 95, 117–127 (2015).

42. Lu, K., Pérez-Gil, J.,(BBA), H. T.-B. et B. A. & 2009, U. Kinematic viscosity of therapeutic pulmonary surfactants with added polymers. Biochim. Biophys. Acta (BBA)-Biomembranes 1788, 632–637 (2009).

43. Kumah, C., Zhang, N., Raji, R. K. & Pan, R. Color Measurement of Segmented Printed Fabric Patterns in Lab Color Space from RGB Digital Images. J. Text. Sci. Technol. 05, 1–18 (2019).

44. Lachmann, B., Robertson, B. & Vogel, J. In Vivo Lung Lavage as an Experimental Model of the Respiratory Distress Syndrome. Acta Anaesthesiol. Scand. 24, 231–236 (1980).

