## Supplementary figures for "Liquid foam therapy (LiFT) for homogenous distribution of exogenous pulmonary surfactant in ARDS"

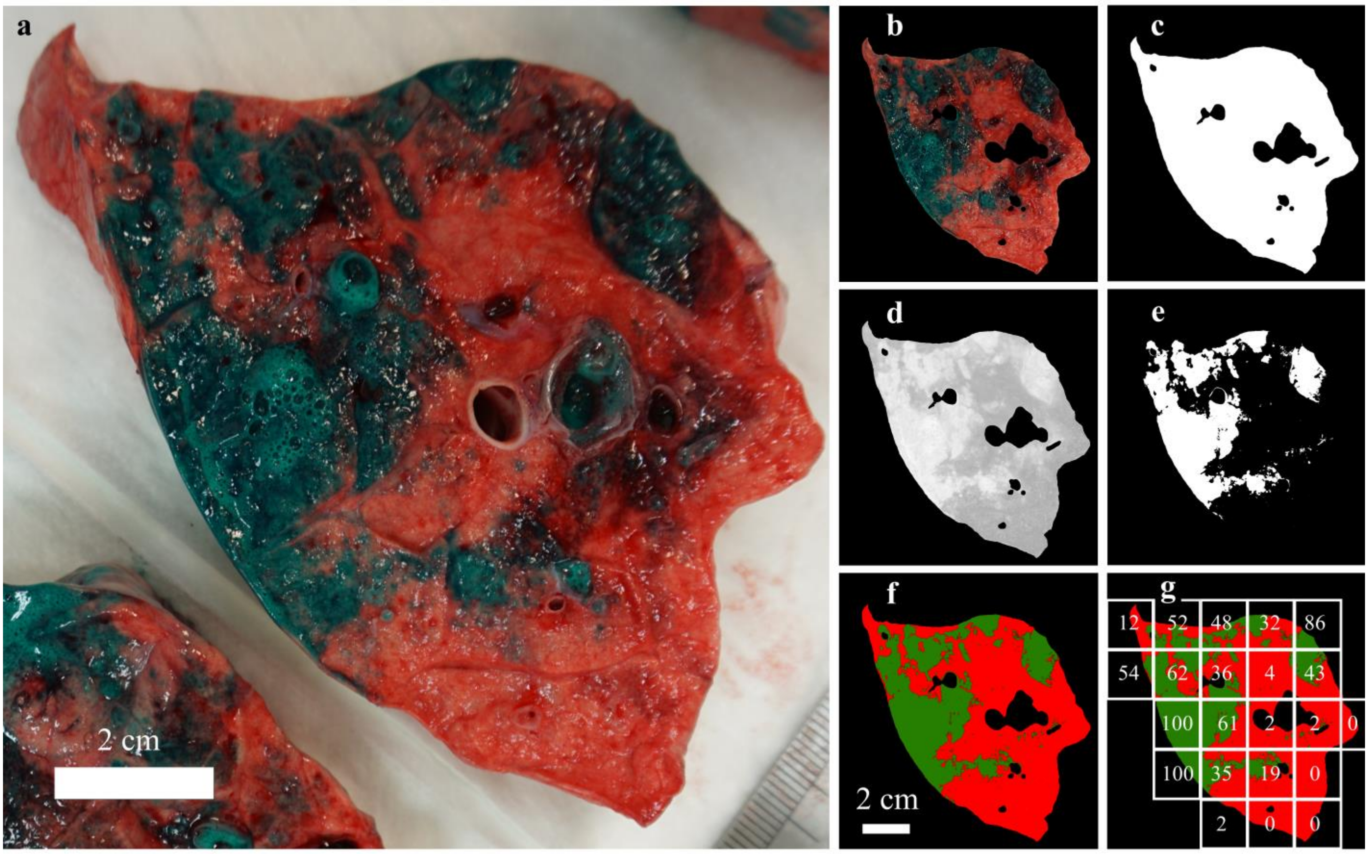


**Figure S1 | Image analysis of lung slices.** (**a**) Example raw image: a cross-sectional cut in the left caudal lobe following *ex vivo* administration of liquid DLSS (distance from caudal edge: *D*=15 cm, same image as in Fig. 1b). (**b-g**) Detailed pipeline of image analysis steps. A masked image (**b**) is obtained from the raw image (**a**) by masking out large cartilage regions, the lumen of large airways and background outside the lung piece. This image is binarized to obtain a ‘mask’ image (**c** and red overlay in **f).** Next, the color scheme of the masked image (**b**) is transformed from RGB to Lab, and the ‘a’ channel (**d**) of the Lab image is thresholded to obtain a ‘dye’ image (**e** and green overlay in **f).** The dyed fraction in the lung slice is calculated as the fraction of white pixels in the ‘dye’ image from white pixels in the ‘mask’ image (34% in the shown slice). The ‘dye’ and ‘mask’ images are then divided into 2×2 cm macropixels (**g**), where the region of the lung slice within a macropixel is referred to as a sub-region. Numbers inside the macropixels correspond to the dyed fraction (%) for each subregion. These values were used to obtain the histogram in Fig. 2b (see main manuscript), showing the normalized number of sub-regions for each dyed fraction interval weighted by the area of the sub-region.

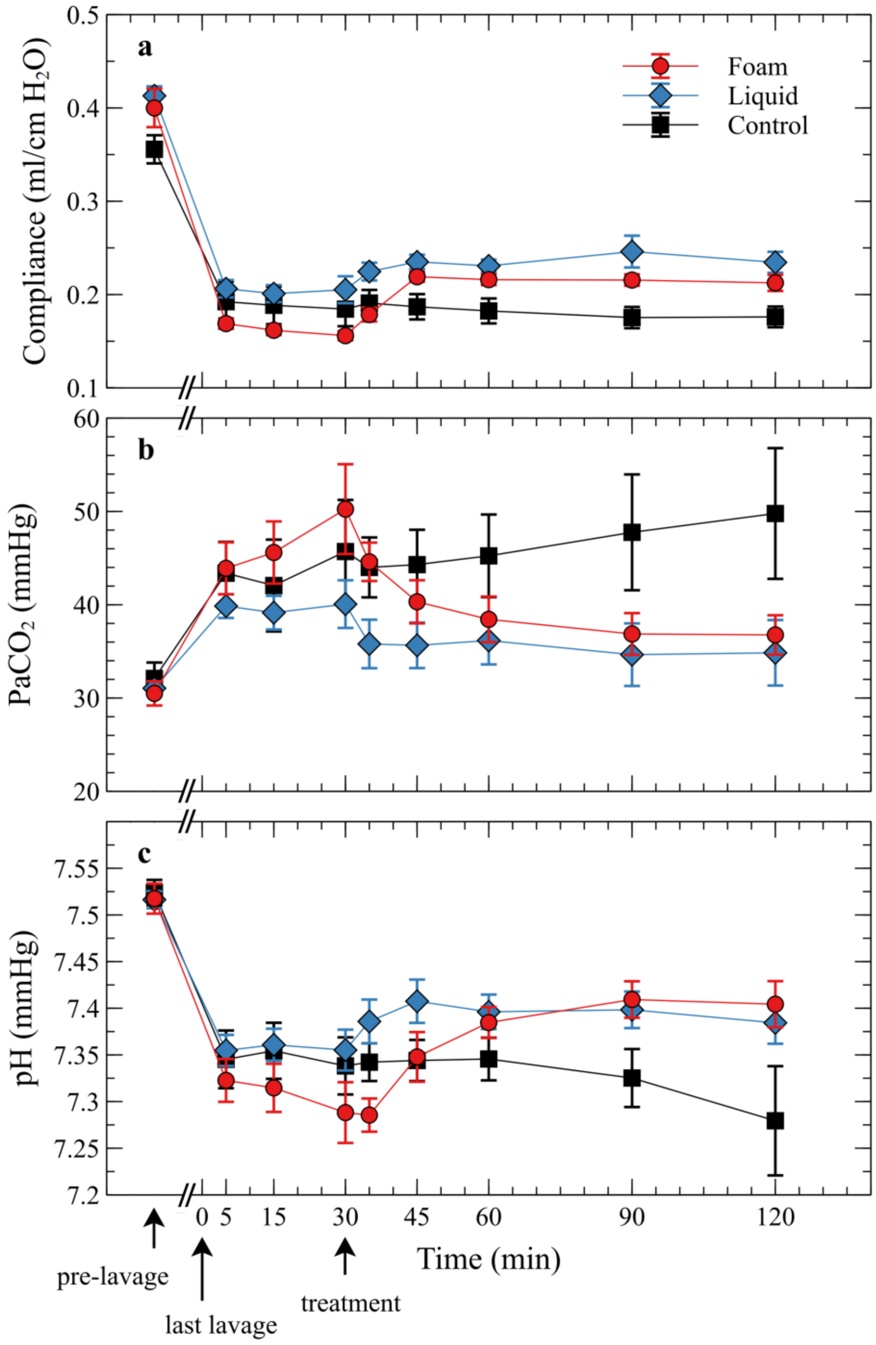


**Figure S2 | Additional *in vivo* results in rats.** Mean (±SE) values measured in the foam, liquid and control groups for (**a**) dynamic compliance (defined as tidal volume divided by the difference between PIP and PEEP), (**b**) arterial $PCO_2$ ($PaCO_2$) and (**c**) pH measurements.

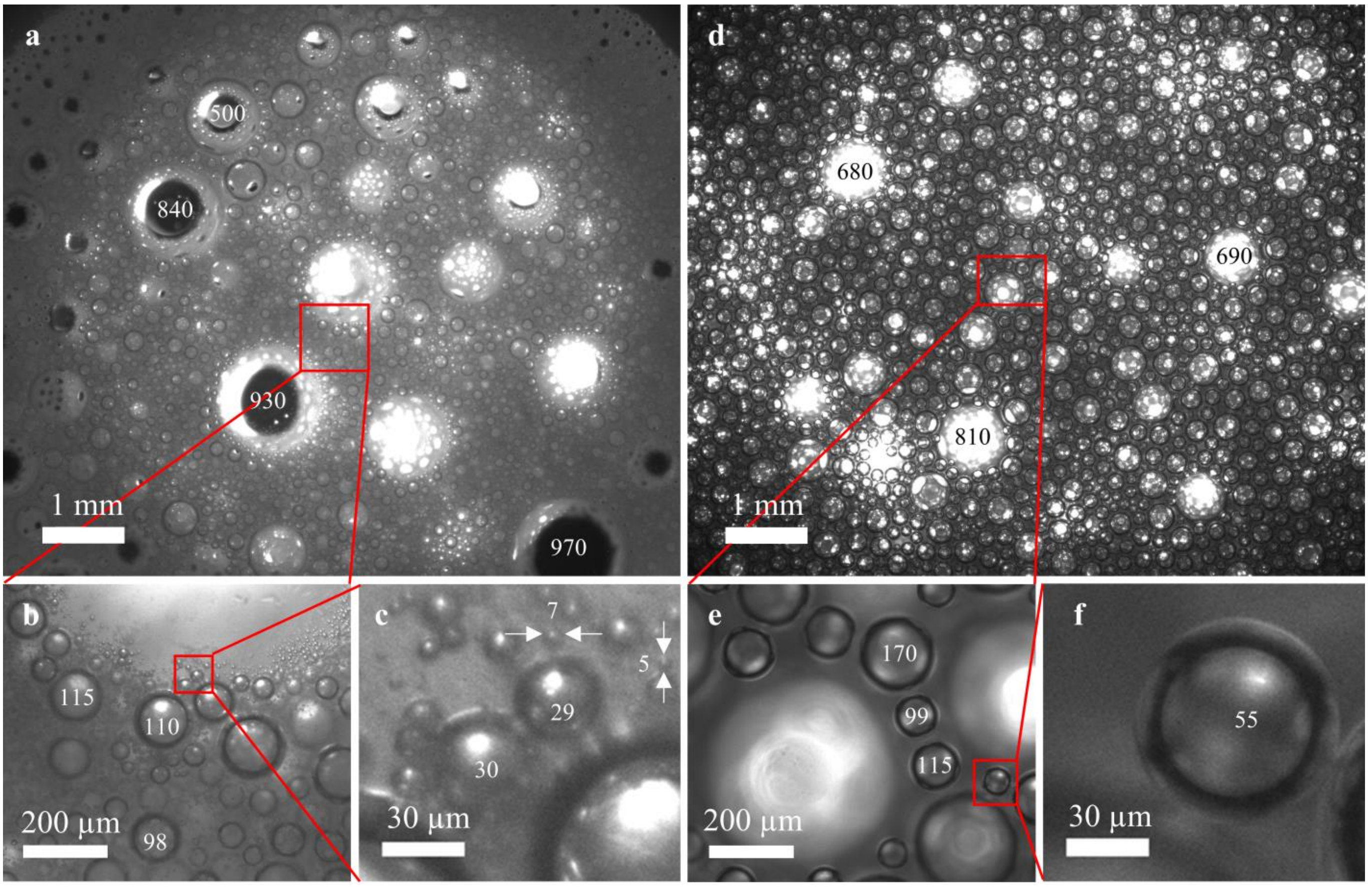


**Figure S3 | Bubble size comparison.** Microscope images of a ~1 mm layer of foamed Infasurf (**a-c)** and foamed DLSS (**d-f**). Images were obtained in bright field mode at 2X (**a,d**) and 10X (**b,c,e,f**) magnification. Numbers inside bubbles represent approximate bubble diameter in micrometers.
